# Fragment-Based Hit Discovery via Unsupervised Learning of Fragment-Protein Complexes

**DOI:** 10.1101/2022.11.21.517375

**Authors:** William McCorkindale, Ivan Ahel, Haim Barr, Galen J. Correy, James S. Fraser, Nir London, Marion Schuller, Khriesto Shurrush, Alpha A. Lee

## Abstract

The process of finding molecules that bind to a target protein is a challenging first step in drug discovery. Crystallographic fragment screening is a strategy based on elucidating binding modes of small polar compounds and then building potency by expanding or merging them. Recent advances in high-throughput crystallography enable screening of large fragment libraries, reading out dense ensembles of fragments spanning the binding site. However, fragments typically have low affinity thus the road to potency is often long and fraught with false starts. Here, we take advantage of high-throughput crystallography to reframe fragment-based hit discovery as a denoising problem – identifying significant pharmacophore distributions from a fragment ensemble amid noise due to weak binders – and employ an unsupervised machine learning method to tackle this problem. Our method screens potential molecules by evaluating whether they recapitulate those fragment-derived pharmacophore distributions. We retrospectively validated our approach on an open science campaign against SARS-CoV-2 main protease (Mpro), showing that our method can distinguish active compounds from inactive ones using only structural data of fragment-protein complexes, without any activity data. Further, we prospectively found novel hits for Mpro and the Mac1 domain of SARS-CoV-2 non-structural protein 3. More broadly, our results demonstrate how unsupervised machine learning helps interpret high throughput crystallography data to rapidly discover of potent chemical modulators of protein function.

## Introduction

Hit detection is a key step in the early stages of the drug discovery process following the identification of a biological target of interest.^1^ A ‘hit’ compound acts as the starting point for the drug design process where the chemical structure of the hit is progressively optimised towards a candidate drug. Approaches towards hit detection generally involve screening large libraries of compounds, both experimentally and computationally.

One of these methodologies is fragment-based drug design (FBDD). In this approach, very low molecular weight compounds (‘fragments’ with typically less than 18 nonhydrogen atoms^2^) are screened at high concentrations against the target protein with X-ray crystallography. A fragment screening approach is more likely to deliver hits than screening larger drug-like molecules because low molecular complexity compounds are more likely to possess good complementarity with the target protein.^3^ Structures of these fragment-protein complexes can then inspire the design of potent binders, either by expanding a fragment to pick up new intermolecular interactions with active site residues, or merging together different spatially proximal fragments.^4,5^ However, despite showing up in X-ray crystallography, the binding affinity of the fragments themselves is typically low. Therefore, gaining potency by fragment expansion or merging is typically a long journey fraught with false starts.

Recently, advances in X-ray crystallography such as automatic crystal mounting robots, fast detectors, as well as increased accessibility to beamtime are enabling high throughput fragment screens. One can routinely go from screening a small fragment library and detecting a handful of hits, to screening 1000s of fragments with ensembles of 100s of fragments hits spanning the binding site.^6,7^ This substantial increase in data enables a systematic data-driven approach for fragment-based hit discovery.

Our key insight is to reframe fragment-based drug design as signal extraction from noisy data by seeking persistent pharmacophore correlations within a fragment ensemble, rather than looking at individual fragments. This is because a fragment itself has low affinity, thus we need the presence of multiple fragments with the same pharmacophore at a particular region of the binding site to provide statistical confidence.

In this work, we employ unsupervised machine learning to learn the spatial distribution of fragment pharmacophores in the binding site. We then use the trained model as a scoring function for virtual screening, picking out molecules with matching pharmacophores. We will first retrospectively validate our model on a dataset of SARS-CoV-2 main protease (Mpro) ligands from COVID Moonshot.^8^ We then present prospective results on identifying hits against Mpro and the Mac1 domain of SARS-CoV-2 non-structural protein 3 (nsp3-Mac1) by performing a virtually screen a library of 1.4 billion purchasable compounds from EnamineREAL.

## Results

### Unsupervised Learning of Pharmacophore Distributions

To turn fragment hits into a model that predicts whether an unknown ligand will bind potently to the binding site, we employ an interpretation inspired by statistical physics. There are multiple chemical motifs that can engage residues on the binding site. These different modes of engagement can be considered as a statistical distribution. Each interaction between a chemical motif on the fragment and a binding site residue corresponds to an instance of this statistical distribution. We assume that the fragment library broadly covers chemical space, and anticipate that stronger interactions will be sampled and therefore observed more often amongst fragment hits than weaker interactions. Note that an individual fragment is a weak binder – fragment screens are done at a high concentration which forces the equilibrium towards forming fragment-protein complexes enabling detection via crystallography. Therefore, we analyse the statistical distribution of fragment-protein interactions formed by the dense fragment hits, rather than any individual fragment (Figure 1a).

**Figure 1:**
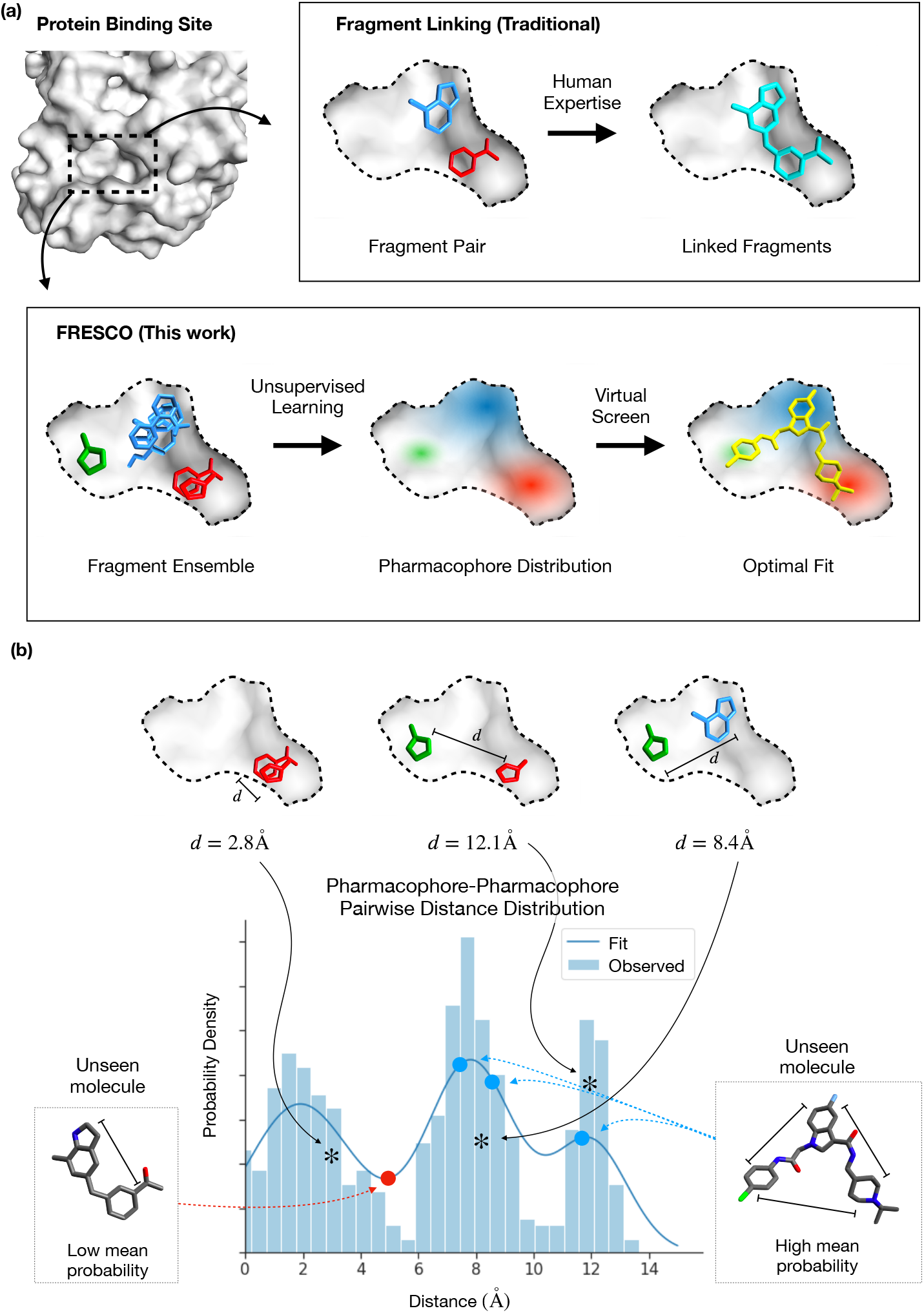
(a) A visual illustration of how FRESCO differs from traditional fragment linking approaches. (b) A visual illustration of how we apply unsupervised learning to fragment ensembles and perform virtual screening of unseen molecules.

To numerically approximate this distribution, we quantify binding interactions by coarse-graining the fragment molecules into hydrogen-bond donor, hydrogen-bond acceptor, and aromatic ring “pharmacophores” (Figure S1). These are a simple abstractions of molecular features that can make potent interactions with binding site residues, and is a commonly used tool to interpret the biological activity of ligands.^9^ The distribution which we then choose to approximate is the pair-wise distance between these pharmacophores. Computational screening of compounds based on pharmacophore distances is a commonly used technique in medicinal chemistry, though here we are extending this concept to enable a statistical interpretation of fragment hit. We consider pharmacophore features, rather than specific protein-ligand interactions, so that the downstream model takes the ligand as the input rather than having to perform the additional step of computationally placing the ligand in the binding site.

We utilise kernel density estimation^10^ to estimate this spatial distribution of pair-wise pharmacophore distances (Figure S2). We then score unseen molecules by evaluating pharmacophore distances within that molecule against the probability distribution of pharmacophore distances derived from the fragment ensemble (Figure 1b). We take the mean probability over all of the distances between all possible pharmacophore-pharmacophore pairs as the score for the molecule. This is an unsupervised approach – starting from the results of a crystallographic fragment screen, without any bioactivity data, we can build a model that computationally screens unseen molecules. We term our approach Fragment Ensemble Scoring (FRESCO).

### Computational Retrospective Study

To validate FRESCO, we perform a retrospective analysis on the COVID Moonshot campaign which is targeting the SARS-CoV-2 main protease (Mpro). Mpro is a target of interest for antiviral drug design as inhibition of Mpro inhibits viral replication, as shown by the recent clinical successes of Paxlovid and Ensitrelvir.^11,12^

The COVID Moonshot consortium^8^ is an open science drug discovery effort which continuously released structures of synthesised molecules along with their bioactivity as the they progressed from initial fragments to optimised leads. This unique dataset allows us to perform a time-split analysis, focusing on the fragment-to-lead phase. We evaluate how our method compares against designs from medicinal chemists, as well as other computational approaches such as docking, and estimate the extent to which FRESCO could have accelerated hit identification.

In the hit identification phase of drug discovery, relatively little is known about what ligand-protein interactions are feasible, thus most proposed molecules are unlikely to be active. A meaningful metric for comparing methods in this regime is the top-*N* “hit rate”, which measures what proportion of the top-*N* predictions by a method are active. We expect the curve from plotting the hit rate against *N* of an informative method to be consistently higher than that of a less informative method. For the Moonshot data we set an IC50 (concentration of inhibitor required to inhibit 50% of protein activity) threshold of 5*μ*M for defining a “hit”. As a point of reference the hit rate in the dataset i.e. the hit rate from compounds designed by expert medicinal chemists, is 6.0%, which illustrates the difficulty of hit detection.

To calculate the hit rate for FRESCO, we first fit a FRESCO model on 23 publicly reported crystallographic structures of non-covalent fragments bound to the SARS-CoV-2 Mpro protein.^7^ We then generate conformers for each Moonshot compound, compute pharmacophore features and pairwise distances, and score the molecules the fitted FRESCO model.

Figure 2 shows that FRESCO achieves higher hit rates compared to both computational docking and the medicinal chemists. Looking at the top-5% of the molecules (*N* < 50), FRESCO has a hit rate of 12-30%, roughly 2-5 times that of the medicinal chemists. Hit rates are also higher than baseline for both lower and higher IC50 thresholds (Figure S3). This shows that it is possible to correlate bioactivity with unsupervised learning of fragment pharmacophore distributions, and that FRESCO could accelerate hit detection in a real-world drug discovery campaign. In this retrospective study, FRESCO is standing on the shoulders of medicinal chemists – it is used to rescore compounds that are designed by chemists. Therefore, we next turn to interrogate the performance of FRESCO when it is used to score a large unbiased library of compounds via a series of prospective studies.

**Figure 2:**
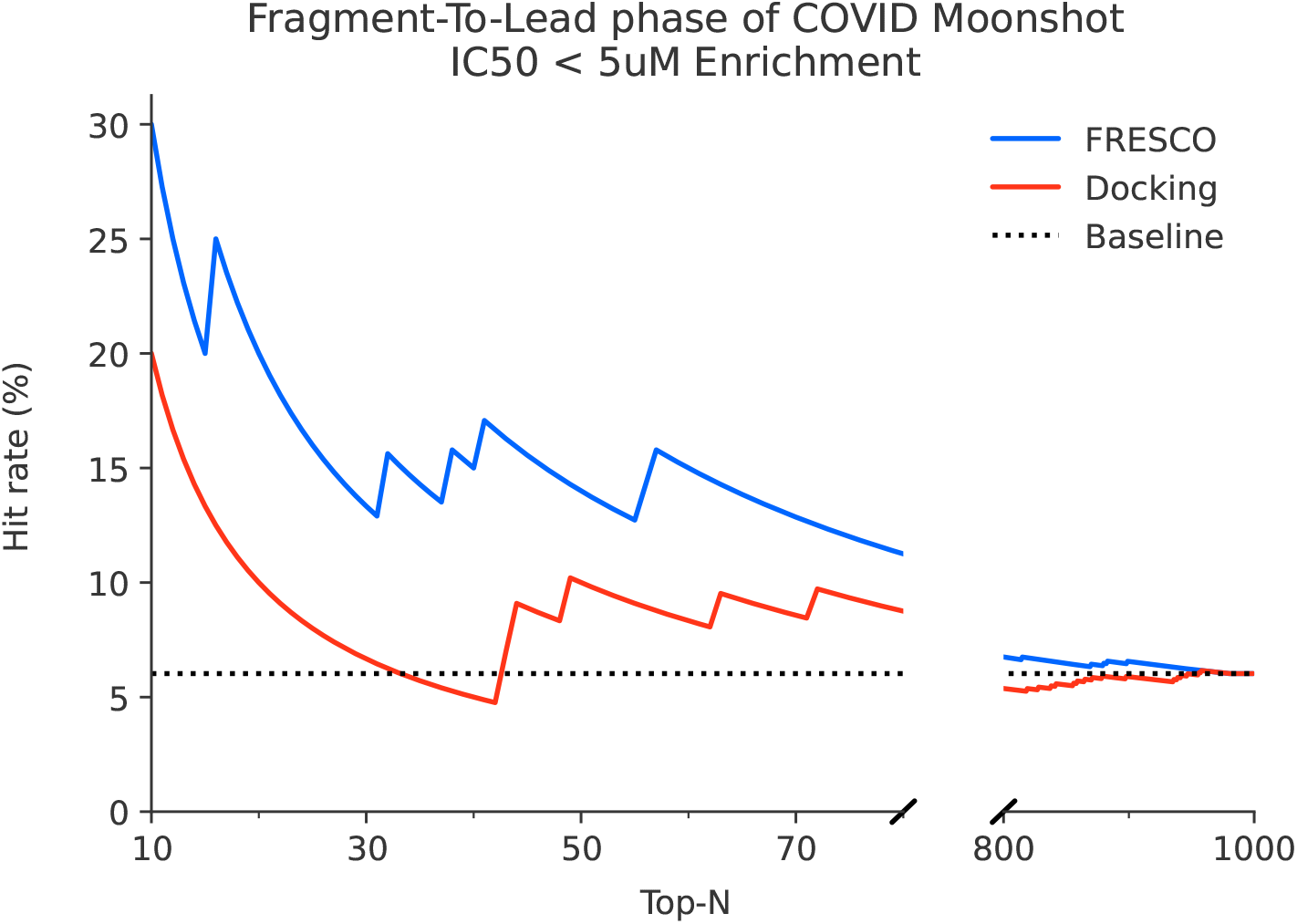
FRESCO is able to retrospectively perform hit detection. High hit rates are achieved relative to docking and the human expert baseline when ranking molecules from the fragment-to-lead phase of COVID Moonshot.

### Hit finding against SARS-CoV-2 Mpro

Building on the results of the retrospective evaluation, we performed a prospective study on Mpro. Rather than rescreening Moonshot compounds, we instead deployed the model to screen the whole Enamine REAL database of 1.4 billion molecules implemented in VirtualFlow^13^ library of commercially available compounds. We then focused the top predicted compounds, filtered them by their physical properties to maximise “drug-likeness”, and selected diverse compounds by clustered hit by structural similarity and picking centroids of the most populous clusters (Figure 3).

**Figure 3:**
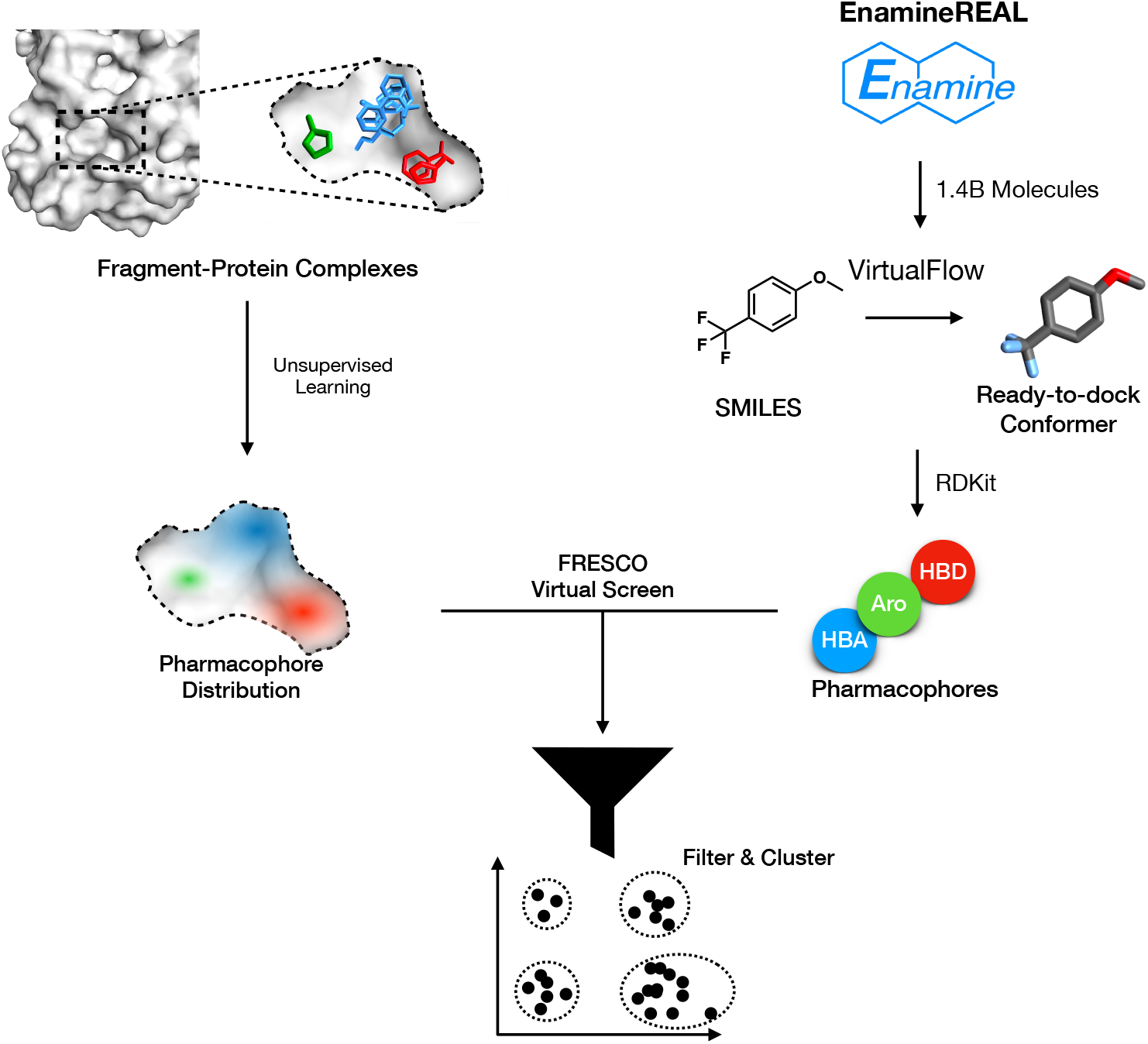
A schematic of the FRESCO screening workflow.

We successfully synthesised and assayed 38 compounds (see SI for the whole library).

The most promising compound, WIL-UNI-d4749f31-37, has an IC50 of 25.8*μ*M measured via fluorescence assay while the remaining compounds were found to be weak-to-negligible activity.

To validate compound activity, we synthesized 8 close analogues to demonstrate the existence of responsive Structure-Activity Relationship^14,15^ (Figure 4). 3 of those compounds, which contained modifications to the 2-hydroxyquinoline substructure of WIL-UNI-d4749f31-37, retained relatively high potency of IC50 < 100*μ*M with one of them (ALP-UNI-ed5cdfd2-1) exhibiting a lower IC50 of 19.4*μ*M. The remaining 5 compounds which perturbed the benzimidazole functional group of WIL-UNI-d4749f31-3 exhibit decreased potency, with only 20-50% inhibition at a concentration of 99.5*μ*M.

**Figure 4:**
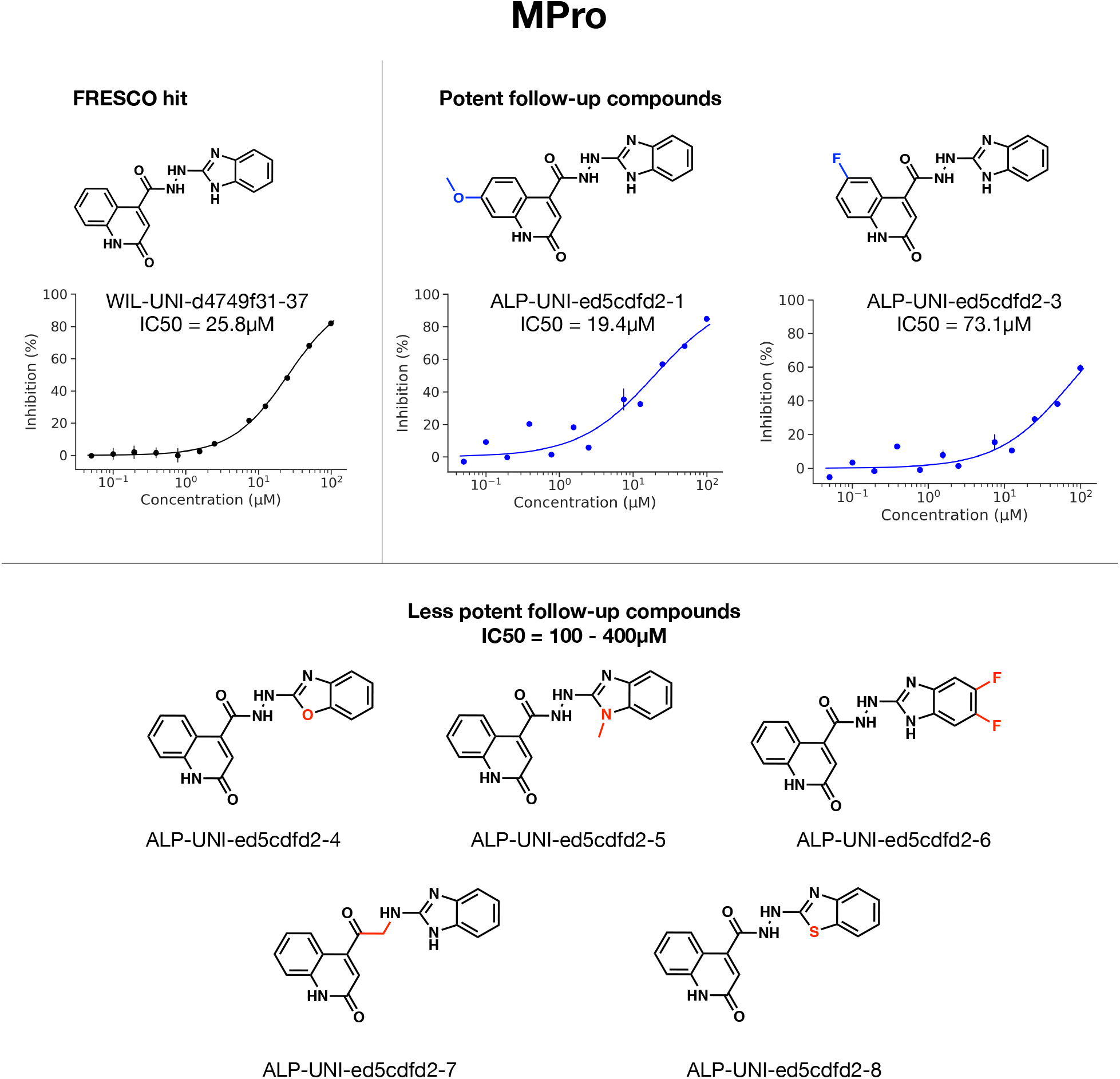
Compound WIL-UNI-d4749f31-37 is identified as a hit against Mpro, with hit confirmation via follow-up compounds demonstrating SAR. Perturbations to the 2-hydroxyquinoline substructure of WIL-UNI-d4749f31-37 led to increased potency while changes to the benzimidazole group consistently decreased potency. Structural differences between the follow-up compounds and WIL-UNI-d4749f31-37 are highlighted in blue/red.

### Hit finding against SARS-CoV-2 nsp3-Mac1

We then turn to SARS-CoV-2 nsp3-Mac1, a structurally unrelated protein target, to demonstrate generalisability of FRESCO in performing hit detection. nsp3-Mac1 is a viral ADP-ribosylhydrolase which counteracts host immune response by cleaving ADP-ribose that is transferred to viral proteins by host ADP-ribosyltransferases. Unlike Mpro, there is no potent chemical matter against nsp3-Mac1. As such, this is a novel first-in-class biological target.

Repeating the FRESCO workflow on a fragment screen against Mac1,^16^ 52 compounds were successfully synthesised and assayed. Two of the compounds show non-negligible activity at high concentration - at 250*μ*M, compound Z5551425673 (as a racemic mixture) has an inhibition of 30.1%, while compound Z1102995175 has 24.8%.

In addition, an X-ray crystallographic screen was also run on the compounds revealing the structure of Z5551425673 (as the S-stereoisomer) bound to the active site (Figure 5). Crystal structures of 9 other compounds chosen via the FRESCO workflow were also obtained though they did not show notable inhibition via HTRF assay (Figure S4). The orthogonal experimental assay and crystal structure results confirm that Z5551425673 is a hit.

**Figure 5:**
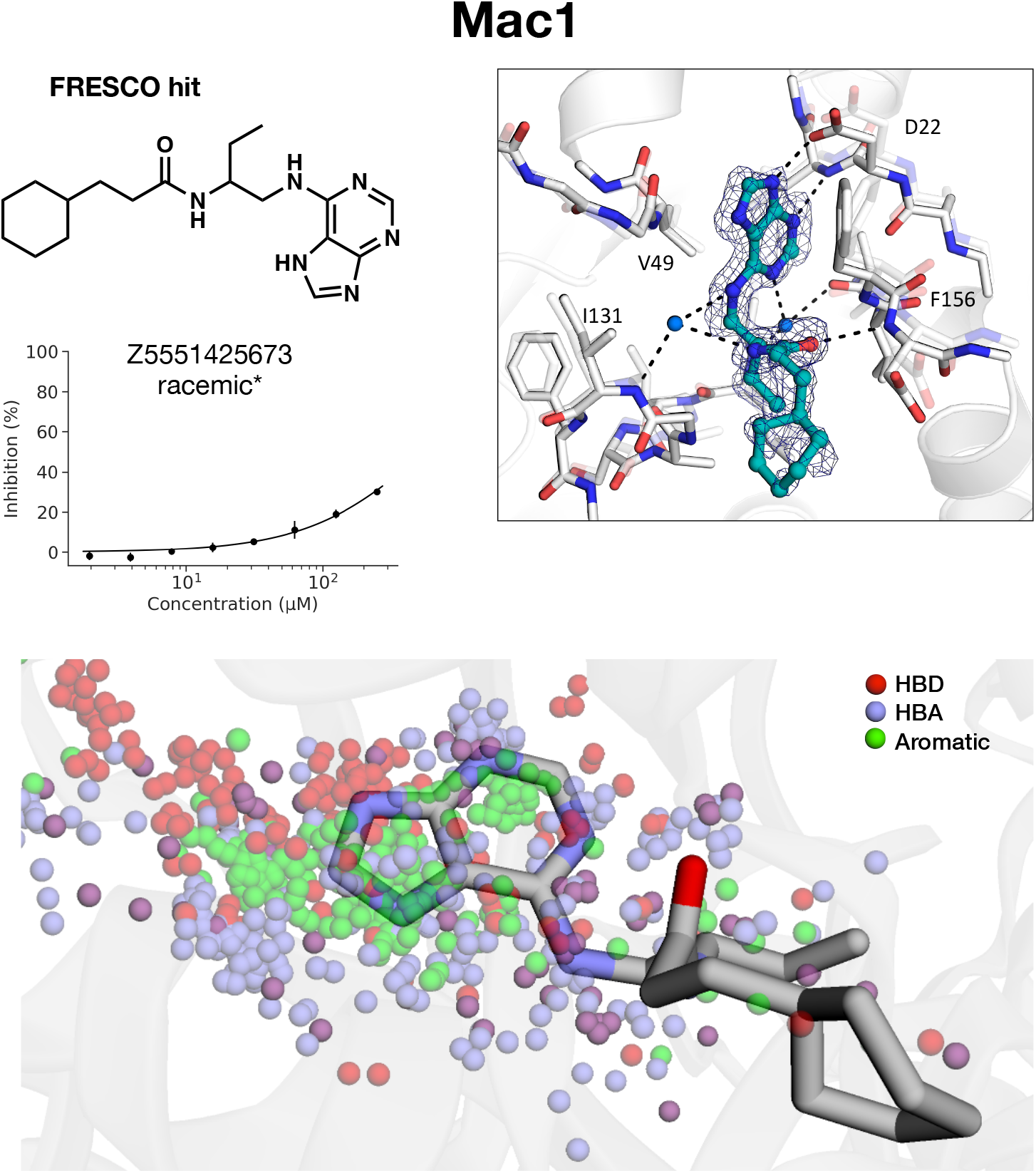
(a) Compound Z5551425673 is identified as a hit against Mac1 via HTRF assay, with (b) hit confirmation via resolution of a crystal structure of Z5551425673 (colored in cyan) bound to the Mac1 active site. (c) The pharmacophores of Z5551425673 match those exhibited by the fragment hits as highlighted by overlaying the bound structure of Z5551425673 (PDB 7FR2) on the distribution of pharmacophores from the fragment ensemble. Note that some functional groups can be regarded as both hydrogen-bond acceptor (blue) and hydrogen-bond donor (red) pharmacophores and hence they are illustrated as purple.

As with Mpro, 11 close analogues to Z5551425673 were ordered to explore the structureactivity relationship of the hit and ensure that the compound is not a singleton. 4 compounds perturbing the aliphatic tail substructure had relatively negligible effect while the remaining compounds perturbing the purine group led to a large drop in activity (Figure 6). These sets of molecules, still weak in potency, are potentially promising starting points for a hit expansion campaign.

**Figure 6:**
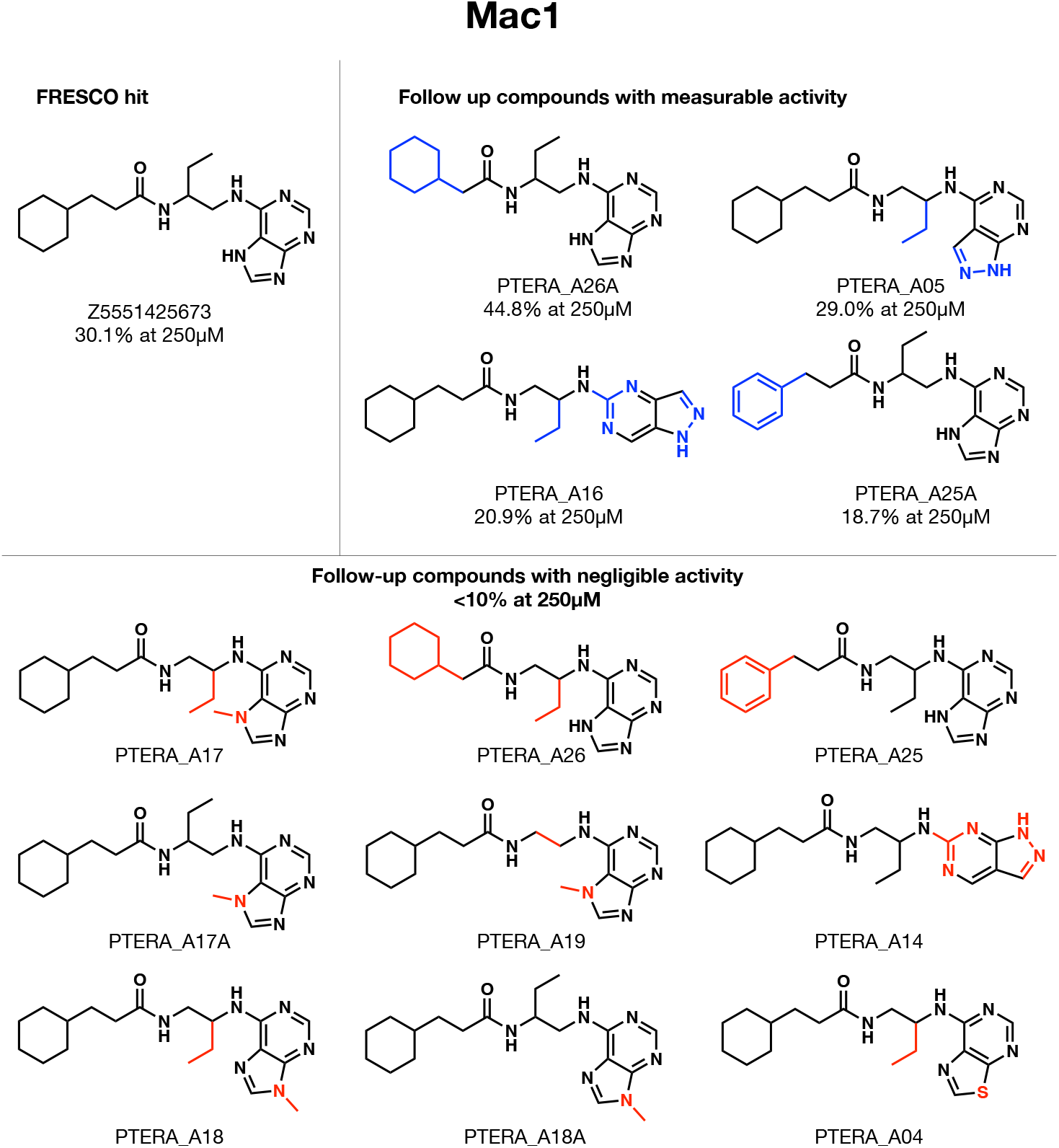
Close analogues around the hit compound identified by FRESCO, Z5551425673, reveals structure-activity relationship which derisks singleton artefacts.

## Discussion and Conclusion

Here we show that the combination of computational statistics with high-throughput structural biology and large libraries of purchasable fragment-like molecules unlocks a powerful tool in hit discovery. Going beyond classical fragment-based drug design, which involves merging or expanding a small set of fragments, we derived a statistical framework that leverages dense fragment hits to build potent inhibitors. Whilst individual fragments are weak binders, our key insight is that a fragment-protein interaction is likely to be significant if there are multiple fragments making similar interactions. Therefore, by picking out these persistent interactions, we can discern the salient chemical motifs which make favourable interactions with the binding site. Specifically, we coarse-grained fragments into pharmacophores, and infer the distribution of pairwise distances between pharmacophores using Kernel Density Estimation. We then screen large libraries of purchasable compounds against this fragment-derived pharmacophore distribution. We retrospectively validated our method using data from The COVID Moonshot, an open science drug discovery campaign against the SARS-CoV-2 main protease, and prospectively discovered new hits against SARS-CoV-2 main protease and nsp3-Mac1.

More generally, we note that our method does not require the observation of affinity data in order to infer potency. This is done by employing an unsupervised machine learning approach on unlabelled structural biology data. As the throughput of structural biology increases, we hope that an unsupervised approach may unlock novel ways of overcoming data limitations in the protein-ligand affinity prediction problem.

## Methods

### Datasets

Fragment crystal structures for model training were downloaded from [Fragalysis](https://fragalysis.diamond.ac.uk/viewer/react/landing). For Mpro, non-covalent fragments from the XChem fragment screen^7^ were used while for Mac1 both XChem and UCSF fragment data were used.^16^

The Moonshot activity data for the retrospective study was accessed in Mar 22nd 2021. The IC50 values in that dataset, as well as in the prospective study on Mpro were measured from a fluorescence based enzyme activity assay, the details of which are described below. To narrow down the data to moleucles during the fragment-to-lead stage of the Moonshot campaign, we only selected molecules which were designed before September 1st, 2020, which gave us a dataset of 979 compounds.

For Virtual Screening, we utilize VirtualFlow, a published dataset of more than 1.4 billion commercially available molecules from EnamineREAL & ZINC15 in a ready-to-dock format.^13^

### Model Construction

The model used in this work takes as input the 3D pharmacophore distribution of a candidate molecule, and evaluates the log-probability that the distribution matches that of the fragment screen on the target site.

The 3D pharmacophore distribution of a molecule is obtained by extracting pharmacophores from the molecular SMILES and their corresponding conformer coordinates, and then evaluating the pairwise distance matrix between all possible pharmacophore pairs (eg Donor-Donor & Aromatic-Acceptor). SMARTS pattern matching following default pharmacophore definitions in [RDKit](https://www.rdkit.org/docs/index.html) were used to extract pharmacophores from the fragment SMILES. The pharmacophores considered are hydrogen bond donors, hydrogen bond acceptors, and aromatic rings. The coordinates of each pharmacophore are defined as the average over the atoms in the pharmacophore (eg the position of an aromatic pharmacophore from a benzene ring would be the mean of the coordinates of the 6 carbon atoms in the ring).

For some fragments, multiple crystallographic poses are recorded. To account for this, we weigh the contribution of each fragment structure to the overall fragment pharmacophore distribution by 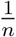 where *n* is the number of conformations recorded for each conformer. In addition, we exclude the counting of correlations between pharmacophores from the same fragment - only correlations between different fragments are measured. This is to avoid spurious intra-fragment correlations that are unrelated to binding to the binding site - strong correlations in pharmacophore distribution between multiple independent fragments are indicative of useful binding interactions and these are what we hope to capture with this methodology.

The bandwidth for KDE fitting was chosen for each system using the Improved Sheather-Jones algorithm^17^ (implemented in [KDEpy](https://kdepy.readthedocs.io/en/latest/index.html)). KDEs of the systems are then constructed using the chosen bandwidths with scikit-learn for technical ease of use in evaluating probabilities. The `scikit-learn` implementation relies on a relatively slow tree-based algorithm that searches over the training datapoints - to increase the efficiency of virtual screening, computationally fast approximations of the KDEs are made using the scipy interp1d function.

Virtual screening of molecular libraries is done by evaluating the probability of the pharmacophore distribution of each molecule using the KDEs in Python. We utilised the VirtualFlow library which provided molecular conformers calculated via ChemAxon and Open-Babel in `PDBQT` format, converted the conformers into `SDF` format, and generated the pharmacophore distributions with `RDKit` as described above. The pharmacophore distributions for each molecule are saved as a pickled dictionary of `numpy` arrays. Each entry of the dictionary contains the flattened pairwise distance distribution for a particular pharmacophore pair. The KDE functions for each pharmacophore combination, trained on the fragment ensemble, are applied onto the dictionary and outputs a mean log-probability for the corresponding pharmacophore combination. The overall score for the molecule is returned as the mean log-probability over all of the pharmacophore combinations.

### Compound Selection

After conducting a virtual screen, the top-500k predictions were selected and filtered to remove undesirable properties. A series of successive filtering steps were performed: first, only molecules with physical properties in well-understood “lead-like” chemical space^18^ were kept. Secondly, the sum of the number of hydrogen bond donors and hydrogen bond acceptors were constrained to an upper limit of 8. Then, we remove molecules that match known filters for pan-assay interference compounds (PAINS)^19^ as well as filters for moieties that are undesirable for medicinal chemistry (eg furan, thiophene, nitro groups). Duplicate tautomers for each molecule are also removed. Finally, for ease of synthetic accessibility, we only consider molecules with less than two chiral centers.

The top-50k molecules remaining from the filtering were then clustered via Butina Clustering^20^ with a Tanimoto distance threshold of 0.2. This resulted in 24748 and 22358 clusters for Mpro and Mac1, respectively. For both targets the centroids of the 50 most populous clusters (or the closest purchasable analogue if it wasn’t available) were chosen as the candidate compounds. These compounds were ordered for synthesis from Enamine which resulted in 38 and 52 successfully made molecules for Mpro and Mac1, respectively.

### Homogeneous Time Resolved Fluorescence assay

The experimental procedure for measuring Mpro inhibition is the same as that previously reported by COVID Moonshot,^8^ which is repeated below.

Dose response assays were performed in 12 point dilutions of 2-fold, typically beginning at 100*μ*M. Highly active compounds were repeated in a similar fashion at lower concentrations beginning at 10*μ*M or 1*μ*M. Reagents for Mpro assay were dispensed into the assay plate in 10*μ*l volumes for a final volume of 20*μ*L.

Final reaction concentrations were 20mM HEPES pH7.3, 1.0mM TCEP, 50mM NaCl, 0.01% Tween-20, 10% glycerol, 5nM Mpro, 37nM fluorogenic peptide sybstrate ([5-FAM]-AVLQSGFR-[Lys(Dabcyl)]-K-amide). Mpro was pre-incubated for 15 minutes at room temperature with compound before addition of substrate and ex/em filter set. Raw data was mapped and normalized to high (Protease with DMSO) and low (No Protease) controls using Genedata Screener software. Normalized data was then uploaded to CDD Vault (Collaborative Drug Discovery). Dose response curves were generated for IC50 using nonlinear regression with the Levenberg-Marquardt algorithm with minimum inhibition = 0% and maximum inhibition = 100%.

Inhibition of SARS-CoV-2 nsp3-Mac1 (aa residues 206-379 of nsp3) was assessed by the displacement of an ADP-ribose conjugated biotin peptide from His6-tagged protein using a HTRF-technology-based screening assay which was performed as previously described.^16^ Compounds were dispensed into ProxiPlate-384 Plus (PerkinElmer) assay plates using an Echo 525 liquid handler (Labcyte). B inding assays were conducted in a final volume of 16 *μ*l with 12.5 nM SARS-CoV-2 nsp3-Mac1 protein, 400 nM peptide ARTK(Bio)QTARK(Aoa-RADP)S (Cambridge Peptides), 1:20000 Anti-His6-Eu3+ cryptate (HTRF donor, PerkinElmer) and 1:125 Streptavidin-XL665 (HTRF acceptor, PerkinElmer) in assay buffer (25 mM HEPES pH 7.0, 20 mM NaCl, 0.05% bovine serum albumin and 0.05% Tween-20). Assay reagents were dispensed manually into plates using a multichannel pipette while macrodomain protein and peptide were first dispensed and incubated for 30 min at room temperature. This was followed by addition of the HTRF reagents and incubation at room temperature for 1 h. Fluorescence was measured using a PHERAstar microplate reader (BMG) using the HTRF module with dual emission protocol (A = excitation of 320 nm, emission of 665 nm, and B = excitation of 320 nm, emission of 620 nm). Raw data were processed to give an HTRF ratio (channel A/B × 10,000), which was used to generate IC50 curves. The IC50 values were determined by nonlinear regression using GraphPad Prism v.9 (GraphPad Software, CA, USA).

### Crystallographic Screening

Crystallographic screening of compounds was performed using Mac1 crystals grown in the P43 space group, following the previously described protocol (PMID: 33853786). Compounds synthesized by Enamine/WuXi were prepared in DMSO to 100 mM and were added to crystallization drops using an Echo 650 liquid handler (Labcyte) (PMID: 28291760). Crystals were soaked at either 10 or 20 mM for 2-4.5 hours, before being vitrified in liquid nitrogen using a Nanuq cryocooling device (Mitegen). Soak times and concentrations are listed in Table S1. Diffraction data were collected at beamlines 12-1 and 12-2 of the Stanford Synchrotron Radiation Lightsource (SSRL). The data collection strategy and statistics are listed in Table S1. Compound binding was detected using the PanDDA algorithm (PMID: 28436492) as described previously (PMID: 35794891). PanDDA was initially run using a background map calculated with 34 datasets collected from crystals soaked only in DMSO (annotated as dmso_34 in Table S1). PanDDA was rerun with a background map calculated using two sets of 35 datasets where no compound binding was detected (annotated as either ssrl_1 or ssrl_2 in Table S1). This procedure led to the identification of an additional nine hits (Table S1).

Compounds were modeled into PanDDA event maps using COOT (PMID: 20383002) with coordinates and restraints generated by phenix.elbow from SMILES strings (PMID: 19770504). Duplicate soaks were performed for most compounds: where the same compound was identified in multiple datasets, the highest occupancy compound was modeled. Both the compound-bound and compound-free coordinates were refined together as a multi-state model following the protocol described previously (PMID: 28436492). Compound occupancy was set based on the background density correction (BDC) value (PMID: 28436492). Refinement statistics are presented in Table S1. Coordinates and structure factor amplitudes have been deposited in the protein data bank (PDB) with the group deposition code G_1002254. PanDDA input and output files have been uploaded to Zenodo (DOI: 10.5281/zenodo.7231822), and the raw diffraction images are available at https://proteindiffraction.org/.

## Acknowledgement

WM acknowledges the support of the Gates Cambridge Trust. AAL acknowledges the Winton Programme for the Physics of Sustainability and the Royal Society. The Mac1 X-ray crystal structures reported in this work were determined using diffraction data collected at the SSRL. Use of the SSRL, SLAC National Accelerator Laboratory, is supported by the U.S. Department of Energy, Office of Science, Office of Basic Energy Sciences under contract no. DE-AC02-76SF00515. The SSRL Structural Molecular Biology Program is supported by the DOE Office of Biological and Environmental Research and by the NIH, National Institute of General Medical Sciences (P30GM133894). NSF Rapid grant 2031205 (to J.S.F.), NIH grant U19AI171110 (to J.S.F.) and TMC Award from the UCSF Program for Breakthrough Biomedical Research, funded in part by the Sandler Foundation (to J.S.F.).

## Supporting Information Available

All code used for this work can be found in the GitHub repo https://github.com/wjm41/frag-pcore-screen. Supplementary figures and tables can be found in an accompanying file.

## Graphical TOC Entry

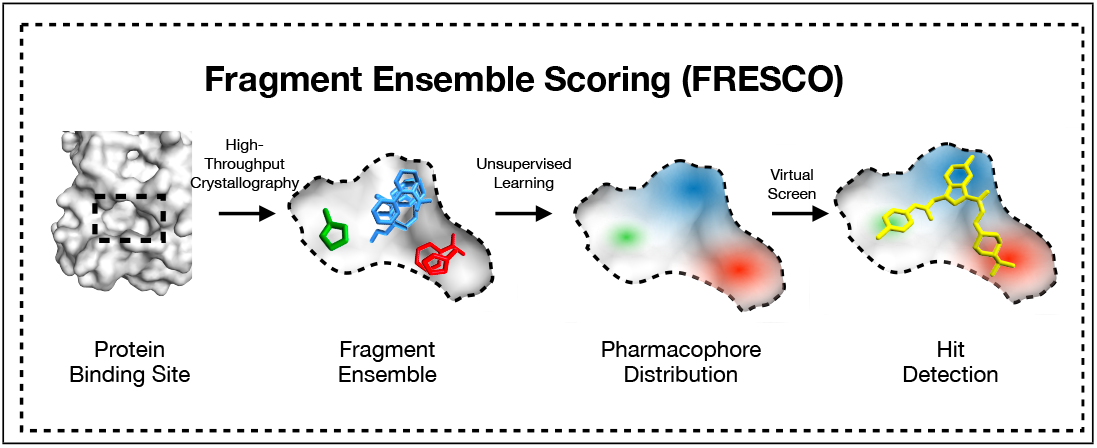

