## Supplementary Information for "Fragment-Based Hit Discovery via Unsupervised Learning of Fragment-Protein Complexes"

#### Supplementary Note 1: Pharmacophore Distributions in 3D

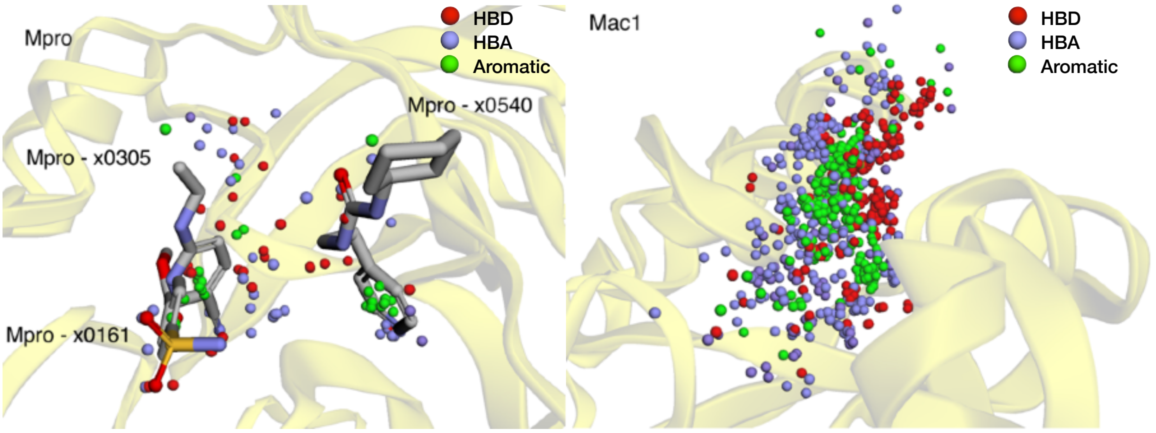


Supplementary Figure 1. The pharmacophores of the fragment ensemble shown in the 3D binding sites of (a) Mpro, and (b) nsp3-Mac1. The red, blue, and green spheres depict hydrogen bond donors, acceptors, and aromatic pharmacophores respectively. Several of the Mpro fragments are drawn to illustrate the 'origin' of some pharmacophores. None are drawn for nsp3-Mac1 due to the density of pharmacophores in the binding site.

### Supplementary Note 2: Pharmacophore Distance Distributions

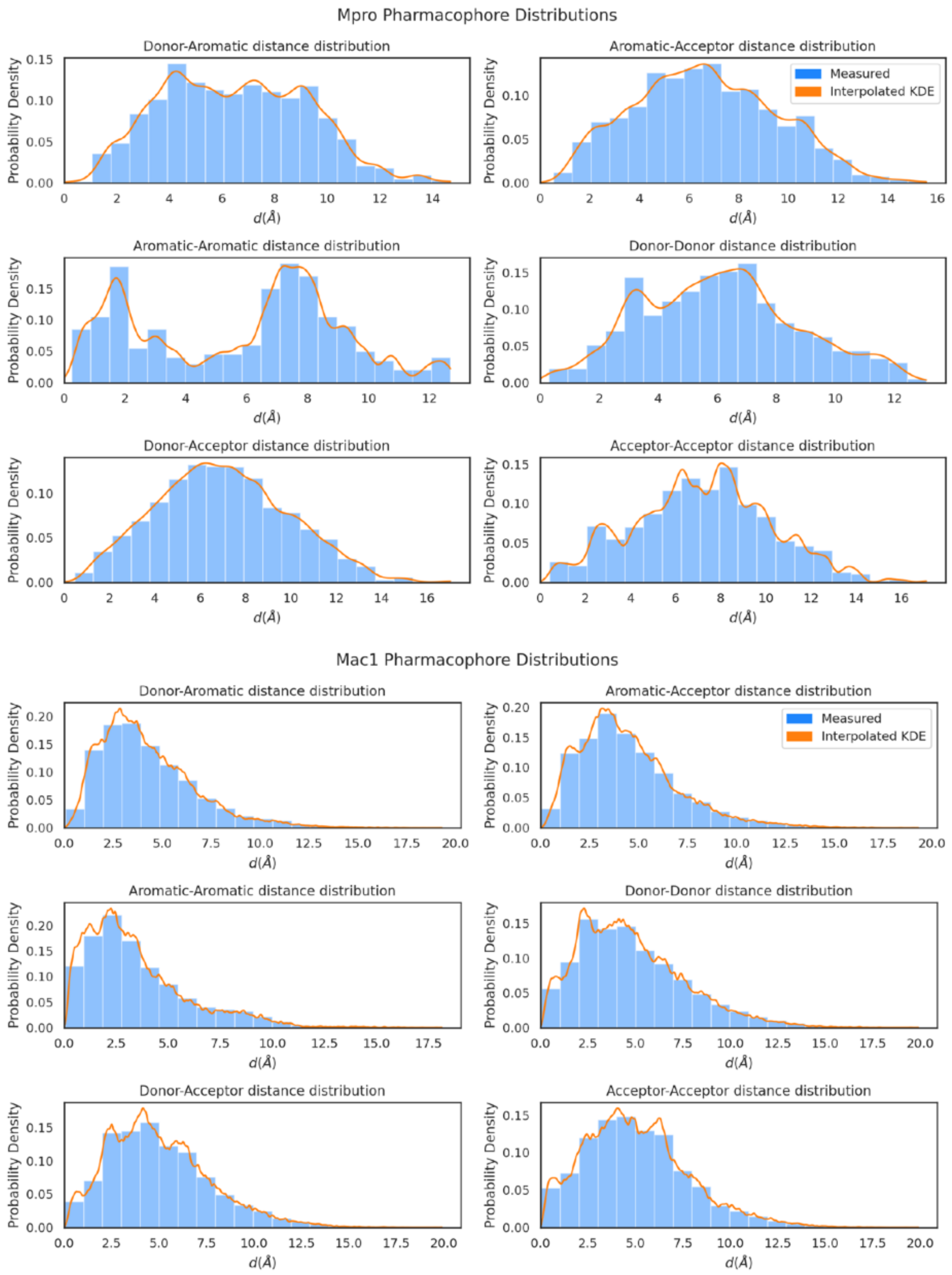


Supplementary Figure 2. The distributions of pharmacophore pairwise distances of all considered pharmacophore pairs for (a) Mpro, and (b) nsp3-Mac1. The histograms in blue are derived by measuring the distribution from the ensemble of experimental fragment-protein complexes, while the orange line shows the KDE fit which is used for scoring.

### Supplementary Note 3: Enrichment plots on COVID Moonshot

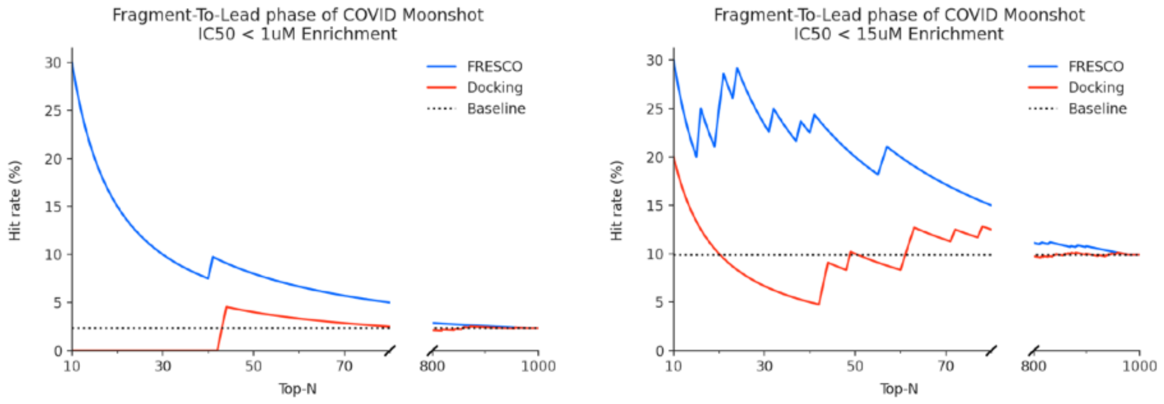


Supplementary Figure 3. The enrichment of FRESCO against Docking on COVID Moonshot activity data as a function of IC50 threshold. The hit rate of all methods increases as the threshold IC50 for defining a ‘hit’ is relaxed.

### Supplementary Note 4: nsp3-Mac1 crystal structures

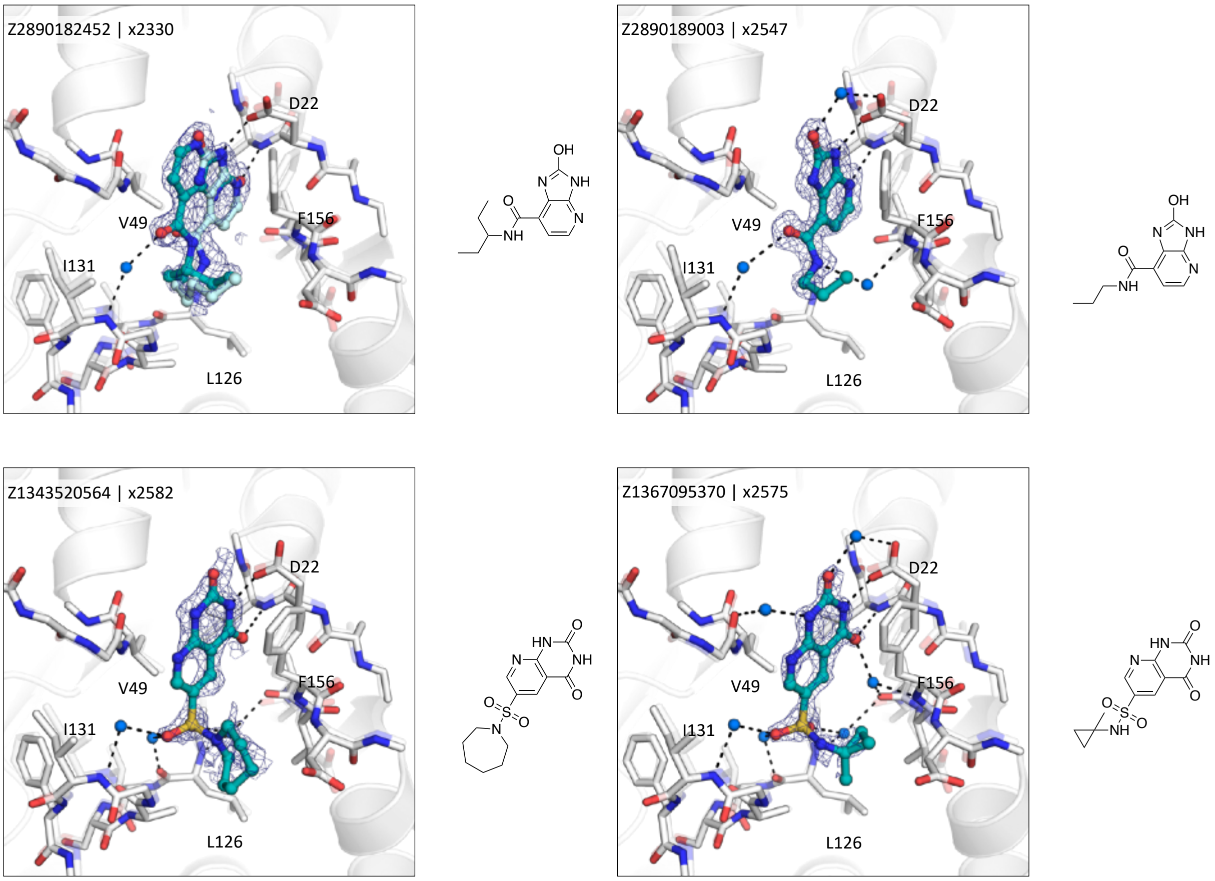

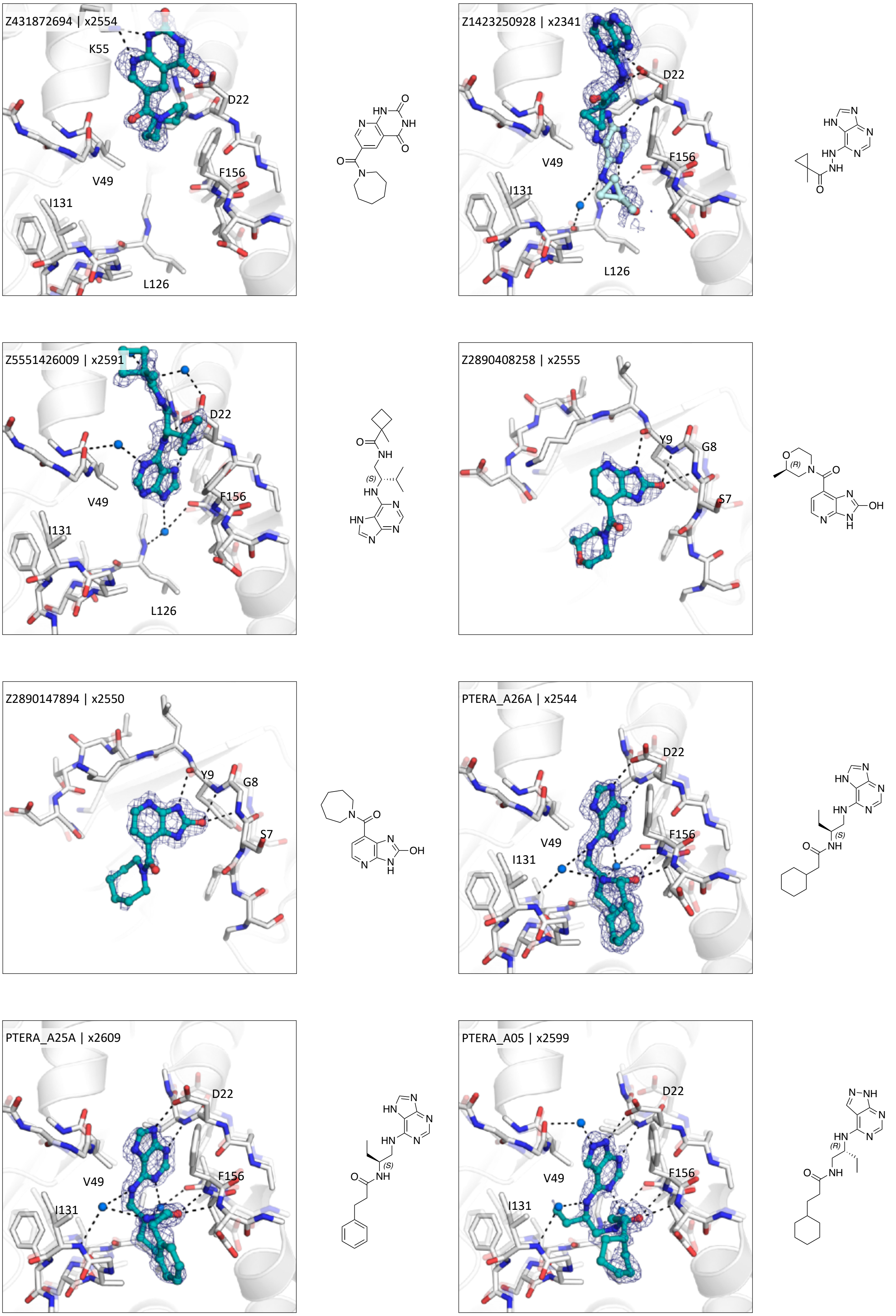


Supplementary Figure 4. nsp3-Mac1 ligand-bound structures obtained from compounds picked by Fresco, as well as follow-up compounds to Z5551425673 (denoted by the PTERA_ prefix) that successfully crystallized. All compounds apart from Z2890408258 and Z2890147894 are crystallised at the active site of nsp3-Mac1. The chemical structure of the compounds are shown on the right of their respective crystal structures.
